# The origin and remolding of genomic islands of differentiation in the European sea bass

**DOI:** 10.1101/223750

**Authors:** Maud Duranton, François Allal, Christelle Fraïsse, Nicolas Bierne, François Bonhomme, Pierre-Alexandre Gagnaire

## Abstract

Speciation is a complex process that leads to the progressive establishment of reproductive isolation barriers between diverging populations. Genome-wide comparisons between closely related species have revealed the existence of heterogeneous divergence patterns, dominated by genomic islands of increased divergence supposed to contain reproductive isolation loci. However, this divergence landscape only provides a static picture of the dynamic process of speciation, during which confounding mechanisms unlinked to speciation can interfere. Here, we used haplotype-resolved whole-genome sequences to identify the mechanisms responsible for the formation of genomic islands between Atlantic and Mediterranean sea bass lineages. We show that genomic islands first emerged in allopatry through the effect of linked selection acting on a heterogeneous recombination landscape. Upon secondary contact, preexisting islands were strongly remolded by differential introgression, revealing variable fitness effects among regions involved in reproductive isolation. Interestingly, we found that divergent regions containing ancient polymorphisms conferred the strongest resistance to introgression.

## Introduction

Understanding how genetic differences accumulate between populations over time to eventually form new species is one of the main objectives in Evolutionary Biology (Mayr 1942; Coyne and Orr 2004). Speciation is generally thought as a gradual mechanism that proceeds through intermediate stages whereby gene flow is not completely interrupted and genomes remain permeable to genetic exchanges (Wu 2001; Feder *et al*. 2012; Harrison and Larson 2014). As long as species can still hybridize, studying gene exchange provides access to the variety of evolutionary mechanisms involved at the different stages of the speciation process (Barton 1979; Hewitt 1988). Advanced sequencing technologies now provide a genome-wide view of divergence between closely related species, improving our understanding of how speciation unfolds at the molecular level (Seehausen *et al*. 2014). A growing number of speciation genomics studies have demonstrated the existence of heterogeneous genomic divergence patterns between entities at different stages of speciation (Turner *et al*., 2005; Harr, 2006; Ellegren *et al*., 2012; Jones et al., 2012; Martin et al. 2013; Gagnaire *et al*., 2013; Renaut et al. 2013; Soria-Carasco et al., 2014). However, difficulties to relate empirical divergence patterns to the underlying mechanisms involved in their formation limit the potential of speciation genomics approaches (Wolf and Ellegren 2016; Ravinet *et al*. 2017).

Heterogeneous genome divergence between taxa can have several possible causes that need to be individually assessed for understanding the underlying mechanisms generating regions of increased divergence (Harrison and Larson 2016; Yeaman *et al*. 2016), the so-called genomic islands (Turner *et al*. 2005; Harr 2006; Nosil *et al*. 2009). Among them, accelerated rates of lineage sorting within populations due to recurrent events of either selective sweeps (Maynard Smith and Haigh 1974) or background selection (Charlesworth *et al*. 1993) can generate incidental islands of relative divergence which are not necessarily related to reproductive isolation (Cruickshank and Hahn 2014). An important objective of speciation research is therefore to identify and understand the origin of genomic islands associated to barrier loci responsible for gene flow reduction between diverging populations (Ravinet *et al*. 2017). Such islands may be themselves explained by different mechanisms depending on the intensity and timing of gene flow, and the genomic architecture of reproductive isolation (Yeaman *et al*. 2016). Elucidating the typical conditions under which each mechanism is at play is central to understanding the roles of selection and gene flow in the speciation process.

The identification of genomic regions that are truly resistant to introgression remains a challenging task, especially because the aforementioned mechanisms are influenced by the recombination landscape and therefore tend to affect similar regions of the genome (Barton and Bengtsson 1986; Nachman and Payseur 2012; Cruickshank and Hahn 2014). To disentangle the role of these confounding factors, substantial levels of gene flow may be needed to properly reveal the genomic regions involved in reproductive isolation. Moreover, the analysis of gene flow and selection may be facilitated by the direct detection of introgressed chromosomal segments, combined with an explicit consideration of the demographic history (Martin and Jiggins 2017). Here, we developed this type of approach in a high gene flow marine species.

The European sea bass (*Dicentrarchus labrax*) provides an interesting model to understand the evolution of genomic islands (Tine *et al*. 2014). The species is subdivided into an Atlantic and a Mediterranean lineage that hybridize in the Alboran Sea (Lemaire *et al*. 2005). Historical demographic inferences revealed that the two lineages have started to diverge in allopatry around 300,000 years BP, and then experienced a post-glacial secondary contact generating varying rates of introgression across the genome (Tine *et al*. 2014). This evolutionary history mirrors the distributional range shifts that occurred across many taxa during glacial periods, which are recognized as an important source of species diversification (Hewitt 1996, 2000). Admittedly, however, the divergence history of sea bass lineages may involve a more complex succession of divergence and contact periods potentially paced by the quasi-100,000-year glacial cycles during the Pleistocene (Snyder 2016). Here, we use haplotype-resolved whole-genome sequences to (*i*) infer the divergence history of sea bass lineages from the length spectrum of introgressed tracts and shared haplotypes, and (*ii*) identify the different mechanisms involved in the formation and remolding of genomics islands of differentiation.

## Materials and Methods

A detailed description of the Materials and Methods is provided in the Supplementary Materials.

### Whole genome resequencing and haplotyping

Haplotype-resolved whole genomes were obtained using a phasing-by-transmission approach (Browning and Browning 2011). Experimental crosses were produced between wild sea bass to generate parent-offspring trios and phase parental genomes using their offspring. Parents were either of Atlantic (ATL, N=4), western Mediterranean (W-MED, N=8) or eastern Mediterranean (E-MED, N=4) origin (Fig. S1). Individual genomes were sequenced to an average depth of 15.5X using Illumina paired-end reads of 100 bp. Reference alignment to the sea bass genome (Tine *et al*. 2014) was performed using BWA-mem (Li 2013). We followed the Genome Analysis Toolkit (GATK) best practice pipeline (McKenna *et al*. 2010; Van der Auwera *et al*. 2013) for variant calling and phasing-by-transmission. After quality filtering to retain only phased variants without missing data, the dataset consisted of 2,628,725 SNPs phased into chromosome-wide haplotypes from 14 individuals.

### Detection of introgressed haplotypes and reconstruction of ancestral Mediterranean genomes

We used Chromopainter (Lawson *et al*. 2012) to identify migrant tracts resulting from introgression between Atlantic and Mediterranean genomes. Chromopainter identifies introgressed haplotype blocks within sequence data by estimating the probability of ancestry from Atlantic and Mediterranean lineages at each variable position along every chromosome using patterns of haplotype similarity. We developed a method to analyze chromosomal ancestral probability profiles in order to determine the starting and ending positions of each migrant tract (Fig. S2). Chromopainter requires non-introgressed reference individuals for every potential source population in order to detect introgressed haplotypes in focal samples. Although the Atlantic lineage is only slightly introgressed by Mediterranean alleles, the Mediterranean lineage is by contrast heavily impacted by gene flow (Tine *et al*. 2014). Therefore, we developed a procedure to reconstruct non-introgressed Mediterranean genomes by removing migrant tracts in a step-wise manner (Fig. S3). Introgressed segments of Atlantic origin were identified by exploiting the different levels of introgression existing between W-MED and E-MED populations. This strategy enabled us to reconstruct the ancestral genetic diversity of the Mediterranean populations before gene flow from the Atlantic, which significantly improved the detection of introgressed tracts (Fig. S4 and S5).

### Analysis of migrant tract length distribution

The distributions of migrant tract length observed in Atlantic and Mediterranean populations were compared to simulated distributions obtained under the secondary contact model previously inferred from the joint allele frequency spectrum (Tine *et al*. 2014). We used the coalescent simulator msprime (Kelleher *et al*. 2016) to generate genome-scale haplotype data with variable recombination rates under this model (Fig. S6 and S7). We then used Chromopainter on simulated data to get the genome-wide distribution of migrant tract length in each population and compare it with the observed distributions.

### Testing waves of historical gene flow

The demographic history of Atlantic and Mediterranean sea bass populations was inferred from the length distribution of tracts of identity-by-state (IBS) using the method developed by Harris and Nielsen (2013). We extended this approach to test for successive waves of gene flow during divergence. We developed a simple and flexible model that can account for multiple equal-length episodes of divergence and gene flow between two populations. Using only 9 parameters, the model can represent a large range of demographic scenarios including (i) continuous migration, (ii) secondary contact and (iii) periodic trains of pulses (Fig. 3A, Fig. S8). The information about the timing of introgression events is expected to be better preserved within low-recombining regions (Pool and Nielsen 2009; Racimo *et al*. 2015). Therefore, we only used sequence information from the low-recombining fraction of the sea bass genome (using *ρ* ≤ 10, the population-scaled recombination rate estimated by Tine *et al*. (2014)) to infer the demographic divergence history from the length distribution of IBS tracts.

### Whole-genome alignment with an outgroup species

We used the 35,012 scaffolds from the *Morone saxatilis* genome assembly to perform alignments against the reference genome of *D. labrax*. The Mauve Contig Mover tool (Rissman *et al*. 2009) from the Mauve software (Darling *et al*. 2004) and the program Abacas (Assefa *et al*. 2009) were used for scaffold ordering and anchoring. Insertions specific to *Morone saxatilis* were removed from the alignment to generate pseudo-chromosome sequences of similar lengths to *D. labrax* chromosomes.

### Population genomics statistics

We used several complementary approaches to assess the level of divergence and introgression between lineages. First, the extent of genetic differentiation between Atlantic and Mediterranean populations of *D. labrax* was evaluated using both relative (*F*_ST_) and absolute (*d*_XY_) measures of divergence. We used Vcftools (Danecek *et al*. 2011) and MVFTools (Pease and Rosenzweig 2015) to calculate the average *F*_ST_ and *d*_XY_ in non-overlapping 100kb windows. Secondly, we used Chromopainter outputs to directly measure the frequency of introgression as the percentage of positions occupied by migrant tracts per window, and the length of migrant tracts along chromosomes. Finally, we calculated the minimum and maximum pairwise distance between haplotypes sampled from Atlantic and Mediterranean populations relative to divergence to *Morone saxatilis* using *RND*_min_ (Rosenzweig *et al*. 2016) and *RND*_max_ statistics. The *RND*_min_ ratio is sensitive to introgression whereas *RND*_max_ rather reflects the level of maximal absolute divergence in a given genomic region. Both statistics are robust to mutation rate variation.

## Results

### Spatial population structure and admixture

The genetic relationships of the newly sequenced genomes with respect to the range-wide population structure of the European sea bass were evaluated with a PCA including 112 additional individuals genotyped at 13,094 common SNPs (Fig. 1A-B, Supplementary Materials). The main component of genetic variation (axis 1, 43.54% of explained variance) clearly distinguished Atlantic from Mediterranean populations, while the second axis (2.59%) revealed a subtle genetic differentiation between eastern (E-MED) and western (W-MED) Mediterranean basins. Genetic admixture was found to occur along the Algerian coast, which is the principal zone where Atlantic alleles enter the Mediterranean Sea. The resulting inflow of Atlantic alleles within the Mediterranean generates a longitudinal gradient of introgression illustrated by a shift between W-MED and E-MED samples along the first PC axis (Fig. 1B).

**Figure 1.**
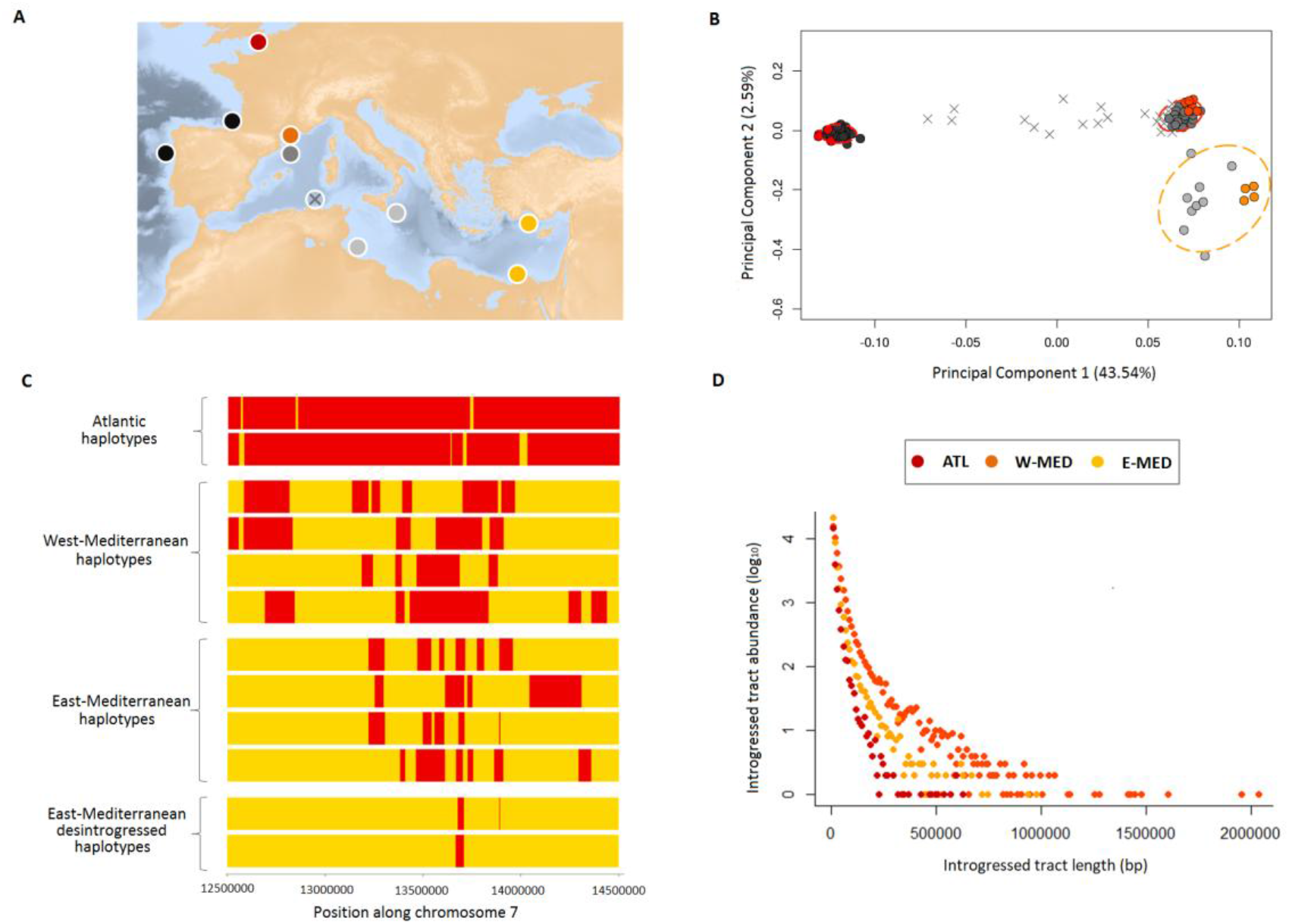
Spatial population structure and introgression. **A.** Geographical location of samples, including the newly sequenced genomes (colored circles) and additional reference samples from the Atlantic (dark grey), W-MED (grey), E-MED (light grey) and the Algerian admixture zone (grey crosses). **B.** Principal Component Analysis of newly sequenced genomes combined with 112 individuals genotyped at 13,094 common SNPs (MAF > 0.1). The first PCA axis distinguishes Atlantic and Mediterranean populations while the second axis reveals a subtle population structure between W-MED and E-MED. Some individuals from the Algerian coast represent admixed genotypes between Atlantic and Mediterranean populations. **C.** Schematic representation of a 2 Mb region within chromosome 7, showing the mosaic of ancestry blocks derived from Atlantic (red) and Mediterranean (yellow) populations. For simplification, we only display two individual haplotypes from Atlantic samples, four from W-MED, four from E-MED, and two “desintrogressed” haplotypes (see text) from E-MED samples. **D.** Migrant tract length distribution obtained for the Atlantic (red, showing tracts of Mediterranean origin), W-MED (orange) and E-MED (yellow) populations (showing tracts of Atlantic origin) using four individuals per population.

### Migrant tracts identification

Spatial introgression patterns also left detectable signatures at the haplotype level along chromosomes (Fig. 1C). The proportion of the genome occupied by migrant tracts was twice higher in the W-MED (31%) compared to the E-MED population (13%), which also displayed shorter migrant tracts (Fig. 1C-D). The longest introgressed haplotype detected in the W-MED (2.03 Mb) was twice longer than the longest one detected in the E-MED population (0.98 Mb). More generally, the genome-wide distribution of migrant tracts length showed a reduced abundance of tracts over all class lengths in the E-MED compared to the W-MED population. This shift is consistent with the action of recombination that progressively erodes recently introgressed tracts as they diffuse by migration from the entrance to the bottom of the Mediterranean Sea. Consistently, this effect was not apparent for the shortest migrant tracts (i.e. < 50 kb) that probably reside in the Mediterranean Sea for a much longer time than the time needed to diffuse from west to east. The Atlantic population was the least introgressed, with less than 5% of its genome occupied by tracts of Mediterranean ancestry. Migrant tracts were also shorter (maximum length 0.62 Mb) and less abundant over the whole length spectrum (Fig. 1C-D). This is consistent with a reduced amount of gene flow from the Mediterranean to the Atlantic population (Tine *et al*. 2014). Finally, our method for reconstructing ancestral Mediterranean genomes effectively removed migrant tracts, since the “desintrogressed” Mediterranean genomes contained very small residual amounts of Atlantic haplotypes (Fig. 1C), except for genomic regions that were strongly introgressed by Atlantic alleles (Fig. S4 and S5, Supplementary Materials).

### Analysis of migrant tract length distribution

An 85% fraction of the 100 kb windows located in low-recombining regions of the Mediterranean genomes present introgressed Atlantic tracts that are on average longer than 50 kb (Fig. S6). Using a recombination clock (Supplementary Materials), we found that this observation is consistent with introgression occurring during the last 17,000 years. Since the most part of the introgressed tracts length distribution is consistent with a recent secondary contact, we evaluated the goodness-of-fit of the previously inferred postglacial secondary contact model (Tine *et al*. 2014), which places the onset of gene flow 11,500 years ago. We performed coalescent simulations with variable recombination rates under this model to generate whole-genome data using the recombination structure of sea bass chromosomes. The length distribution of migrant tracts obtained from these simulations reproduced well the observed distribution for both the Atlantic and Mediterranean populations (Fig. 2). Therefore, the secondary contact model inferred from the joint site frequency spectrum without using linkage information has a high predictive power regarding the length distribution of introgressed tracts when accounting for recombination rate variation.

**Figure 2.**
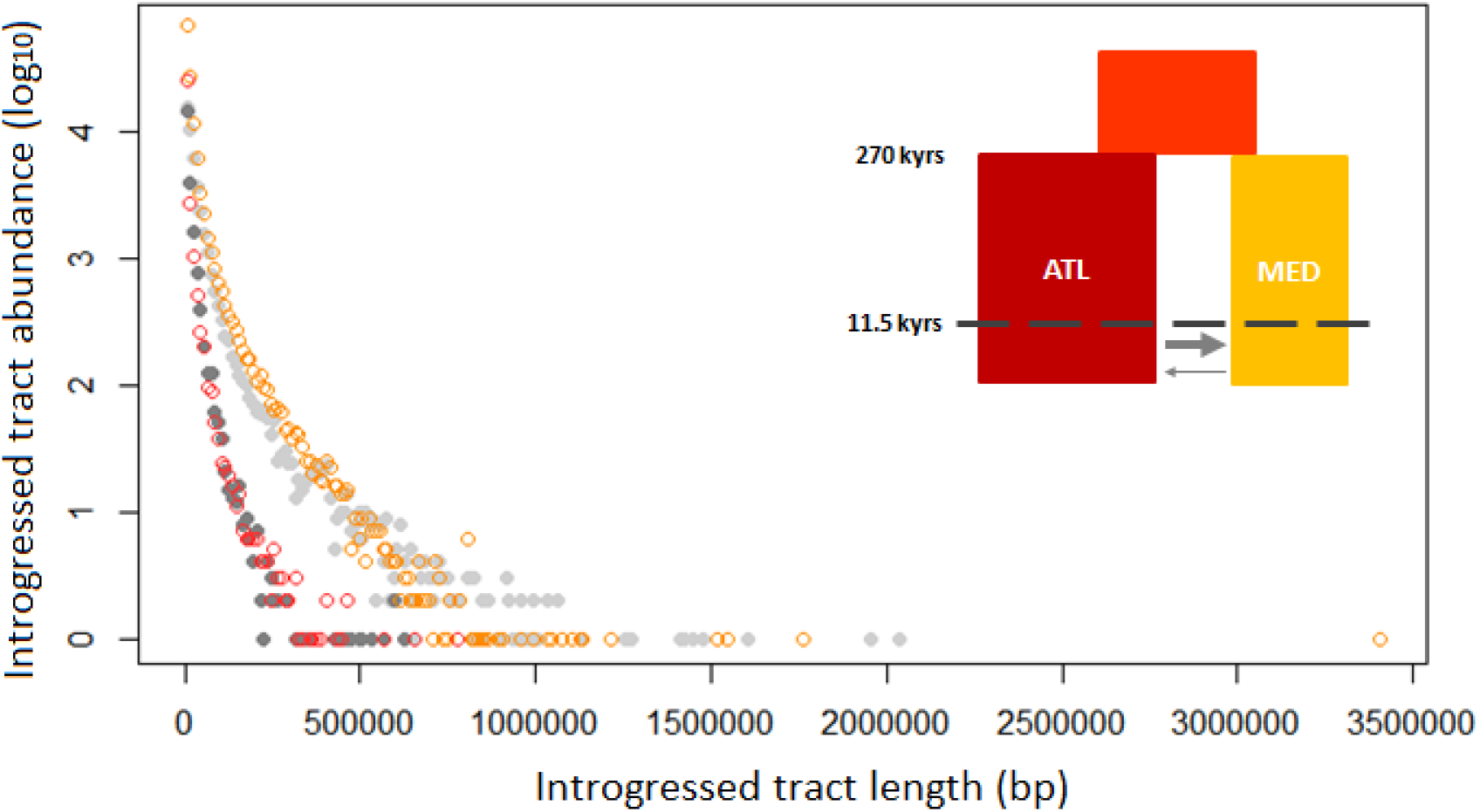
Observed and simulated migrant tract length distributions in the Atlantic and Mediterranean populations. Observed distributions (grey) of migrant tract length are compared with simulated distributions (colored) under the post-glacial secondary contact scenario inferred by Tine *et al*. (2014), illustrated in the top-right corner. The abundance of introgressed tracts as a function of their length is represented for observed vs. simulated data in the Atlantic (dark grey vs. red circles, showing tracts of Mediterranean origin) and Mediterranean populations (light grey vs. yellow circles, showing tracts of Atlantic origin).

### Testing waves of historical gene flow

Initial divergence between sea bass lineages has been dated around 270,000 year BP (Tine *et al*. 2014), corresponding to three glacial cycles (Snyder 2016) during which possible genetic interactions may have occurred when interglacial conditions were similar to present. In order to address whether short migrant tracts found within low-recombining regions of the genome could result from waves of historical gene flow, we developed a flexible model of divergence in which the history of admixture can take different forms (Fig. 3A). The modeling scenarios were subdivided into three categories according to the distribution of admixture pulses over time: (i) continuous migration, (ii) secondary contact, and (iii) periodic trains of pulses (Fig. 3B). Our demographic inferences showed an increase in likelihood with increasing numbers of pulses in each scenario (Fig. 3C). This is because the total number of admixture pulses contained in each model (i.e. the product *m×n*) acts as a hidden nuisance parameter, even though the three different scenarios were built using the same number of model parameters. Therefore, we only compared likelihood values among scenarios with identical number of admixture pulses, considering up to 10 pulses in total (the likelihood tended to flatten out beyond this value). We found the secondary contact scenario to be the best supported model across the entire range of admixture pulse number (Fig. 3C and Table S1), and therefore found no support for a periodic pulse model with separate waves of gene flow. The best-fit secondary contact model (n = 10 pulses, Fig. 3D) provided clear support for asymmetric introgression, with a more than 6-fold higher introgression rate from the Atlantic into the Mediterranean than in the opposite direction, which is consistent with previous findings (Tine *et al*. 2014). The duration of allopatric divergence relative to the secondary contact period was found shorter than previously reported (Table S2). Nevertheless, the splitting times estimated between Atlantic and Mediterranean sea bass lineages were highly consistent across methods (*ca*. 300,000 years BP for the best IBS tract model vs. 270,000 years BP from the best JAFS model (Tine et *al*. 2014)).

**Figure 3.**
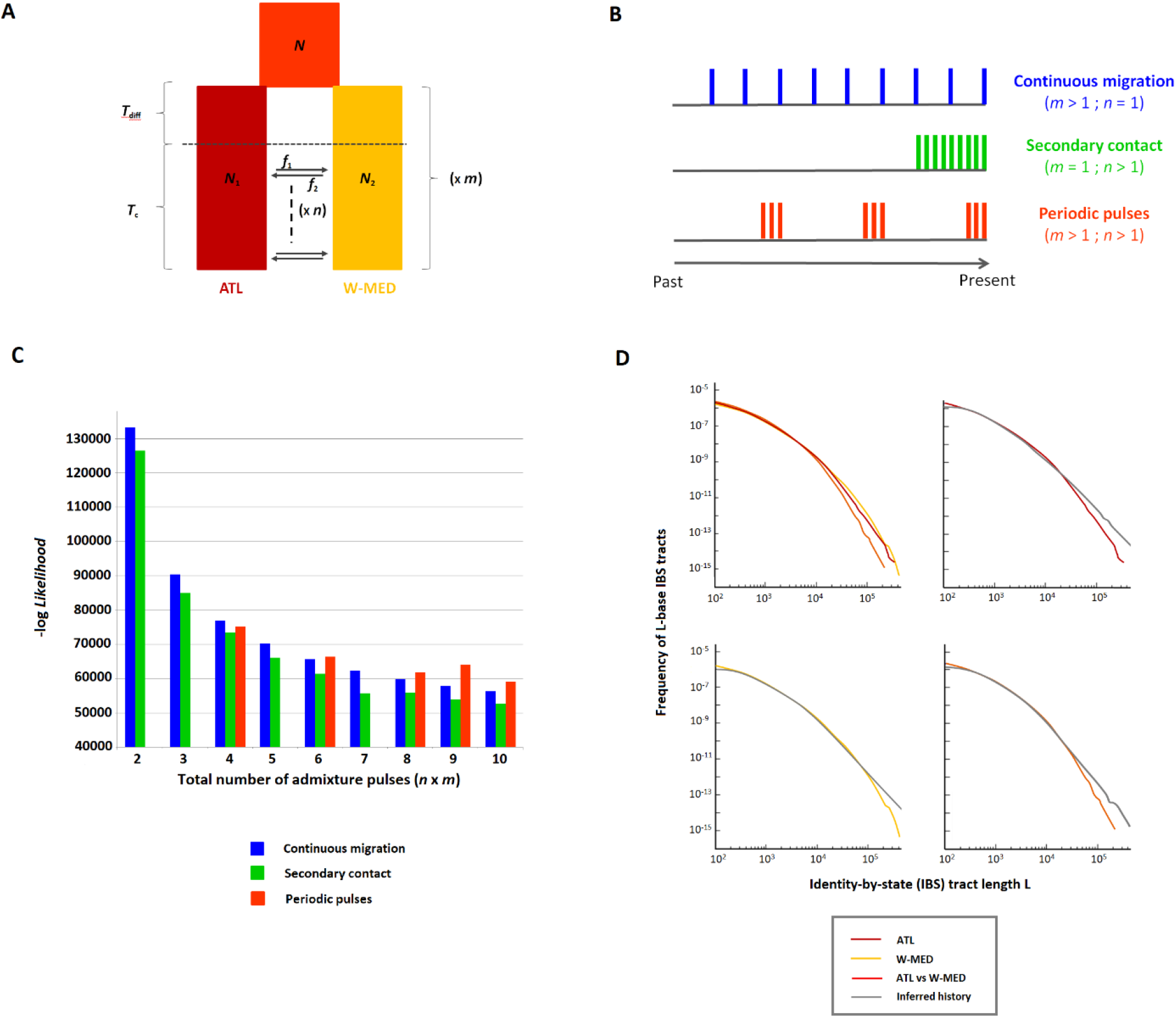
Demographic history inferred from the length distribution of IBS tracts. **A.** Flexible demographic model accounting for multiple equal-length episodes of divergence and gene flow between Atlantic and Mediterranean sea bass populations. An ancestral population of size *N* splits into two populations of size *N*_1_ and *N*_2_, experiencing one to several (*m*) cycles of interrupted gene flow during *T*_diff_ generations followed by migration during *T*_c_ generations. Each contact episode contains one to several (n) pulses of admixture (black arrows), replacing Mediterranean and Atlantic populations by a proportion *f*_1_ and *f*_2_ of migrants, respectively. The *n* admixture pulses are homogeneously distributed along each contact episode. **B.** Modeling scenarios fall into three different categories according to the distribution of admixture pulses over time: continuous migration, secondary contact, and periodic trains of pulses. The illustrated example shows these three categories with 9 pulses represented by vertical bars along the timeline. **C.** The log-likelihood values obtained for the different categories from two to 10 admixture pulses. Two possible configurations of the periodic pulses scenario exist for models including a total of 6, 8 and 10 pulses (Table S1), only the best of which is represented here. The periodic pulses scenario is not defined for 2, 3, 5 and 7 pulses. **D.** Goodness-of-fit of the best model (*m*=1, *n*=10), showing the length distributions of IBS tracts from observed data (colored lines) compared to model prediction (grey lines). Upper-left: the three distributions observed within ATL, W-MED, and between ATL and W-MED populations. Upper-right: model fit within ATL. Lower-left: model fit within W-MED. Lower-right: model fit between ATL and W-MED.

### Chromosomal profiles of divergence and introgression

We investigated chromosomal patterns of genetic differentiation (*F*_ST_) and absolute sequence divergence (*d*_XY_) between Atlantic and Mediterranean populations. We found highly varying levels of relative and absolute divergence across the genome, with Mb-scale regions of elevated divergence preferentially mapping to low-recombining regions (Fig. 4A-C and S9). Consistent with predictions from the “linked selection hypothesis” (Cruickshank and Hahn 2014; Burri 2017a), *F*_ST_ and *d*_XY_ were respectively negatively and positively related to the population-scaled recombination rate (*ρ* = 4*N*_e_*r*) (Fig. 5A-D, S10 and S11). However, the highest *F*_ST_ values tended to be associated with high values of *d*_XY_ mapping preferentially to low-recombining regions (Fig. 5), which is not expected under the only action of linked selection (Burri 2017b). In order to evaluate the extent to which this observation is consistent with the existence of genomic islands resistant to gene flow (Guerrero and Hahn 2017), we compared divergence with measures of introgression. We found that the most divergent windows in terms of both *F*_ST_ and *d*_XY_ are also the ones that strongly resist gene flow, either detected by high *RND*_min_ values (>0.05, Fig. 5E-F and S12) or low frequencies of introgressed tracts (<0.1, Fig. 5G-H and S13). This is confirmed by the similar levels of ancestral and contemporary *F*_ST_ values in these regions (Fig. 4A-B, mauve vs. purple *F*_ST_ plots in the red dotted box). Therefore, barriers to gene flow between Atlantic and Mediterranean sea bass tend to involve regions of low-recombination where both relative and absolute divergence are higher than expected under the sole effect of linked selection. The value of *F*_ST_ alone is not a good indicator of the barrier strength, since regions with similar *F*_ST_ values display variable levels of resistance to introgression (Fig. 5E-F). This is illustrated by the finding of islands of differentiation that have been partly eroded by gene flow since secondary contact, especially in the W-MED population (Fig. 4A, mauve vs. purple *F*_ST_ plots in the black dotted box).

**Figure 4.**
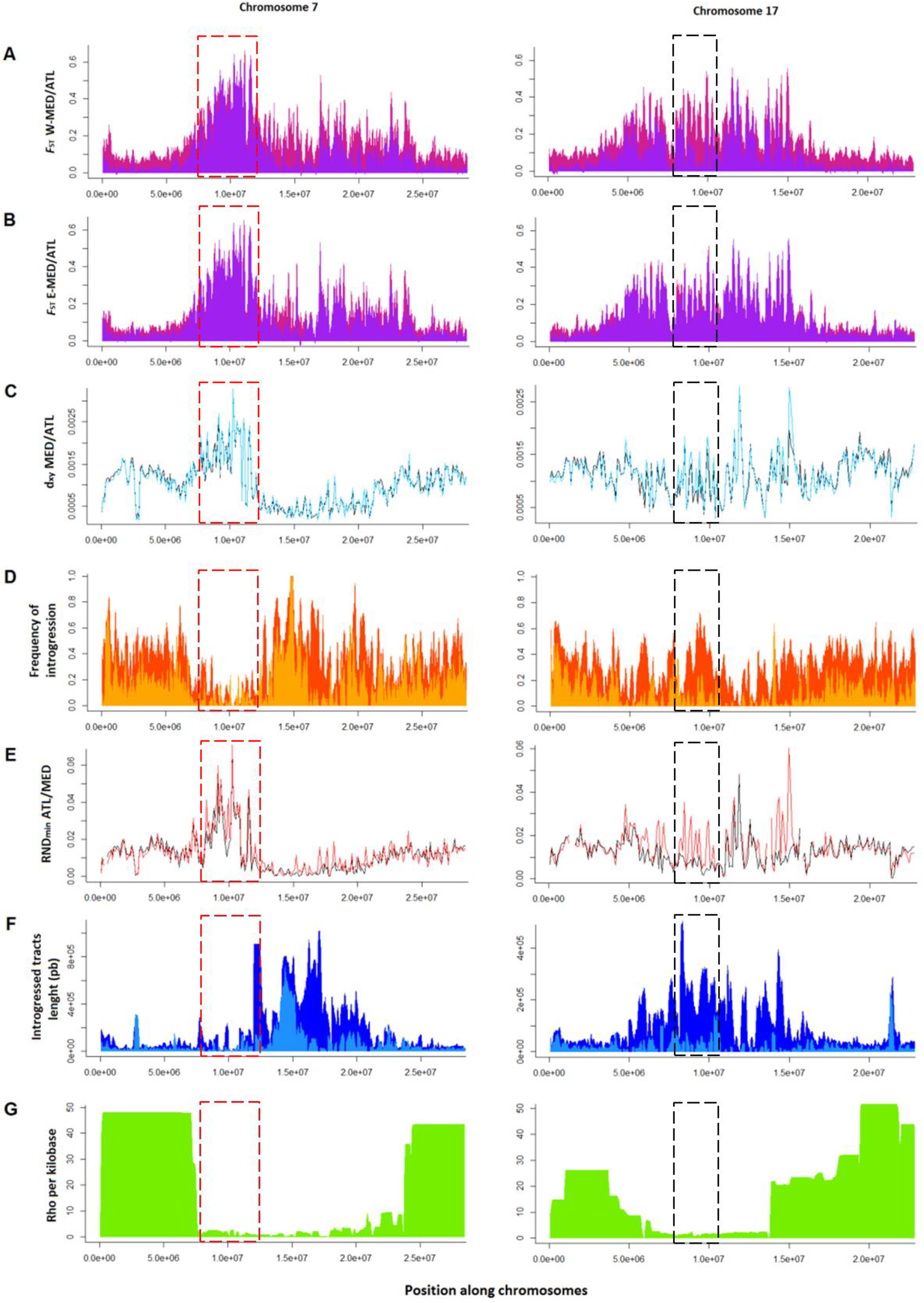
Population genetic statistics calculated in non-overlapping 100 kb windows along chromosomes 7 and 17. **A.** *F*_ST_ measured between the Atlantic and contemporary (purple) or ancestral reconstructed (mauve) W-MED or E-MED (**B.**) population. **C.** *d*_XY_ calculated between the Atlantic and the W-MED (black) or E-MED (blue) population. **D.** Frequency of introgression in the W-MED (orange) or E-MED (yellow) population. **E.** *RND*_min_ measured between the Atlantic and W-MED (black) or E-MED (red) population. **F.** Average length of introgressed tracts in the W-MED (dark blue) or E-MED (light blue) population. **G.** Population-scaled recombination rate (*ρ* = 4*N*_e_*r*) averaged between Atlantic and Mediterranean population, inferred by Tine *et al*.

**Figure 5.**
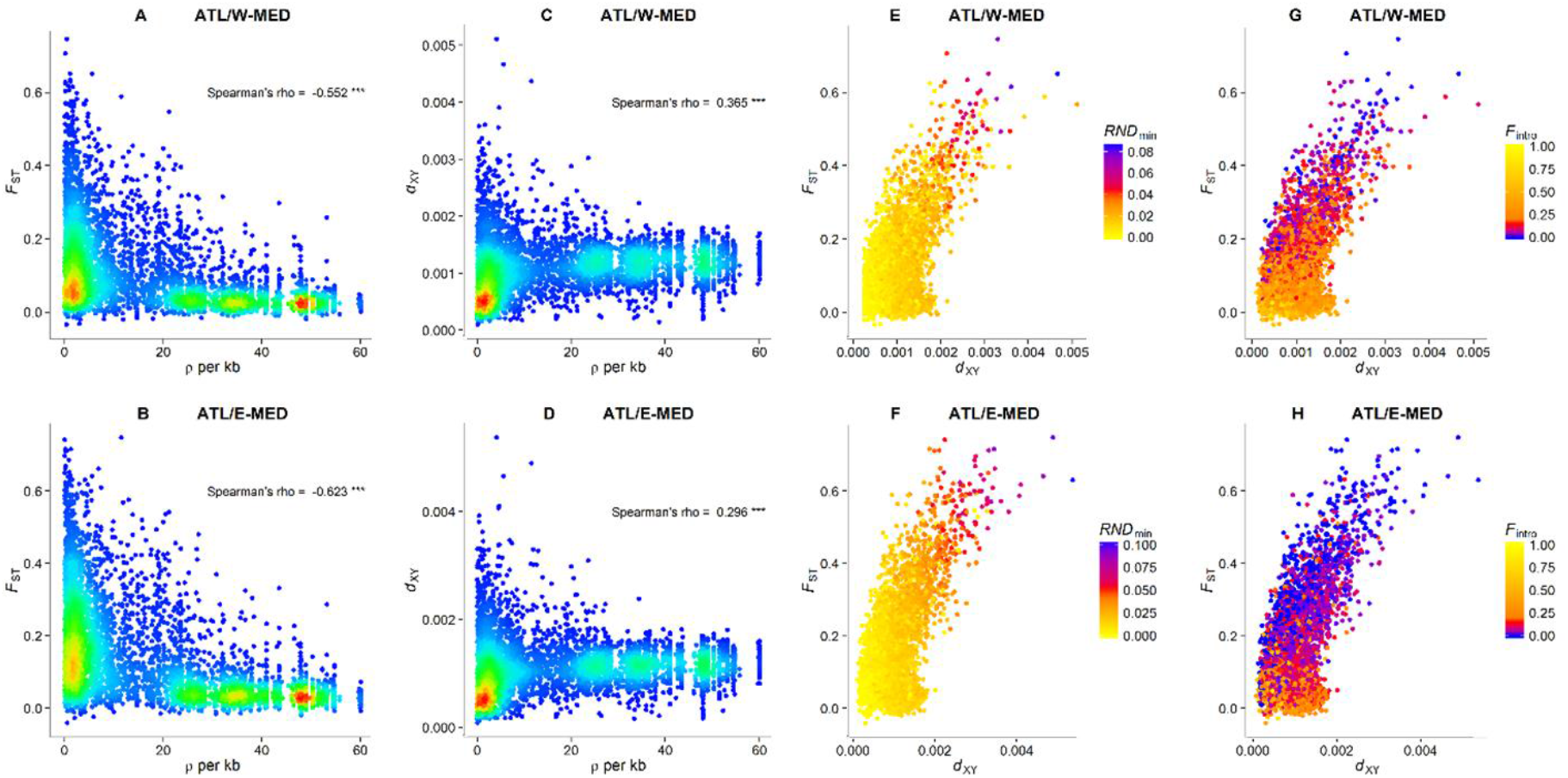
Relationships between divergence (*F*_ST_ and *d*_XY_), the population-scaled recombination rate (*ρ* = 4*N*_e_*r*) and introgression statistics (*RND*_min_ and *F*_intro_). **A-D.** The density of points appears in color scale from low (blue) to high (red) densities. **E-G.** The color scale indicates the value of *RND*_min_ (**E-F**) or the frequency of introgression (**G-H**) in the corresponding window from low (blue) to high (yellow) introgression rate values.

Variable degrees of resistance to gene flow among genomic regions were also detected using spatial comparisons of divergence patterns. Despite the strong correlation observed between ATL/W-MED and ATL/E-MED *F*_ST_ patterns (Fig. S4), some peaks of ancestral differentiation almost completely vanished in ATL/W-MED contemporary patterns, but remained remarkably unchanged in the ATL/E-MED comparison (Fig. 4, black dotted box). By contrast, some peaks seem to have resisted to introgression in both ATL/W-MED and ATL/E-MED comparisons (Fig. 4, red dotted box). These differences are consistent with variable strength of the barrier effect among genomic regions involved in reproductive isolation.

We also tested the theoretical prediction that introgressed tracts closely located to sites where selection against introgression is strong should be shorter compared to the average tract length (Sedghifar *et al*. 2016). At the genome scale, the length of introgressed tracts was largely determined by the local recombination rate, and tracts at a given position were on average longer in the W-MED compared to the E-MED population (Fig. 4F-G). However, by combining information across the strongest barriers to gene flow identified with outlier values of *RND*_min_, we found that the mean length of introgressed tracts was reduced (up to four-fold) in the close vicinity (< 200 kb) of the barrier loci (Fig. S14), consistent with theoretical predictions.

Finally, in order to determine if the strength of barriers to gene flow is related to the degree of past differentiation and maximal divergence between Atlantic and Mediterranean alleles, we evaluated the effect of these two factors on the frequency of introgression. Both ancestral *F*_ST_ and *RND*_max_ were found to negatively affect the frequency of introgression (Fig. 6). This was not a methodological artifact, since our desintrogression method would on the contrary tend to overestimate the ancestral *F*_ST_ in highly introgressed regions, where the reconstructed ancestral diversity of the Mediterranean population may be downwardly biased. Therefore, our results indicate that the regions with the highest level of pre-contact differentiation and maximal divergence were the less likely to introgress during the recent secondary contact episode.

**Figure 6.**
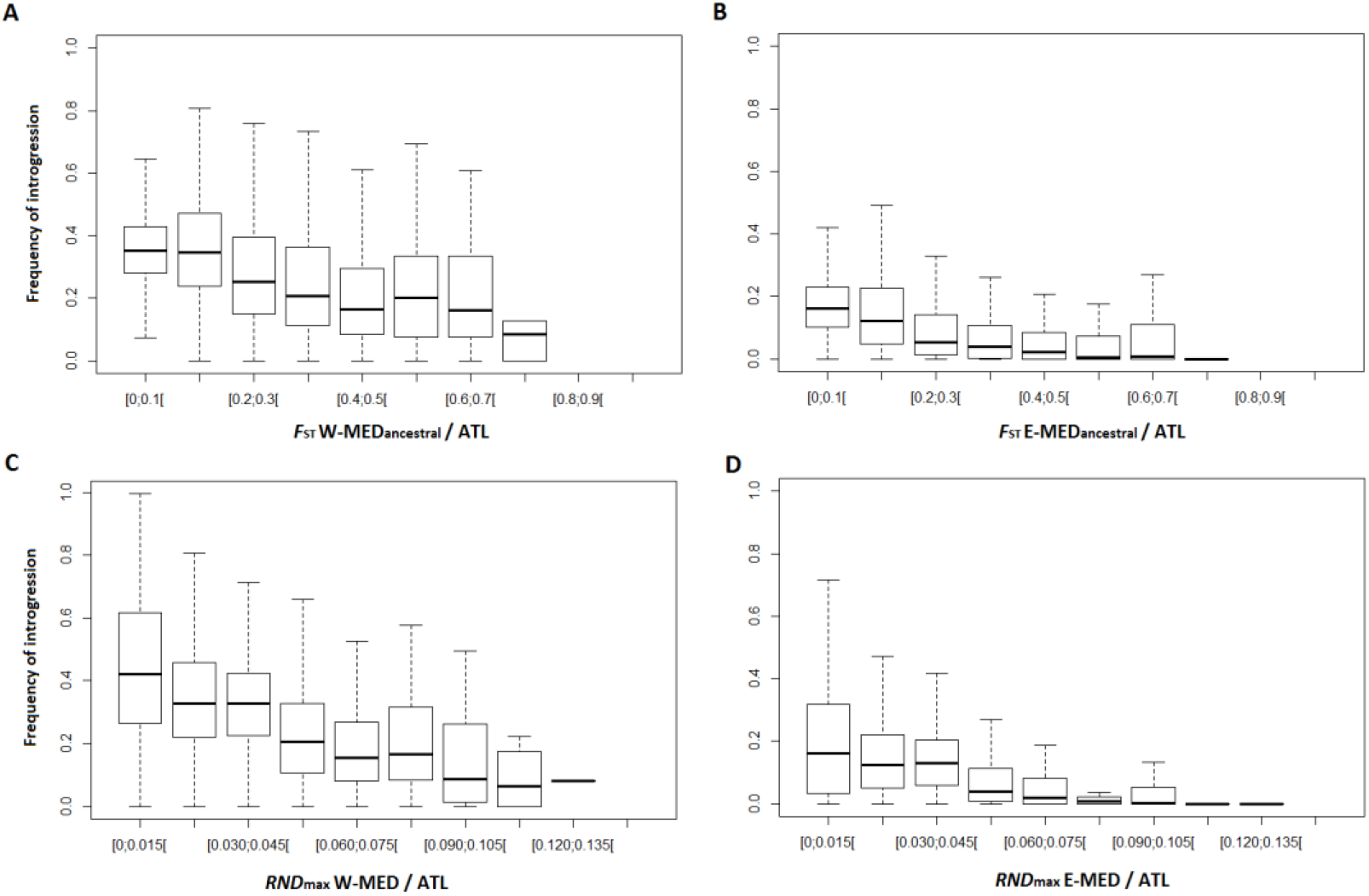
Frequency of introgression for different levels of past genetic differentiation and allelic divergence measured with *RND*_max_. **A.** Introgression as a function of past *F*_ST_ measured between the Atlantic and the reconstructed ancestral W-MED or E-MED (**B.**) population. **C.** Introgression as a function of *RND*_max_ measured between the Atlantic and W-MED or E-MED (**D.**) population. Each box represents the lower and upper quartiles and the median of introgression frequency values.

## Discussion

### Inference of divergence history from phased data

We used two very different approaches based on haplotype information to resolve the history of divergence and gene flow between Atlantic and Mediterranean sea bass lineages. First, our coalescent simulations with recombination showed that the observed length distribution of introgressed tracts can be well reproduced by the postglacial secondary contact model inferred in a previous study (Tine *et al*. 2014). We then evaluated whether the presence of short introgressed tracts in low-recombining regions could indicate the existence of older admixture events. Although there is a possibility that during the whole divergence history, the quasi-100,000-year glacial cycles (Snyder 2016) have promoted discrete waves of gene flow, we did not find evidence for a cyclic connectivity model representing glacial oscillations. Instead, our demographic inferences based on the IBS tract spectrum also supported a scenario of secondary contact and confirmed that Atlantic and Mediterranean sea bass lineages have started to diverge around 300,000 years BP.

The only disagreement between the two approaches concerned the relative duration of the divergence and contact periods. Admittedly, the power of our inferences may be limited by the confounding effects of time and recombination on the length of introgressed tracts (Pool and Nielsen 2009; Liang and Nielsen 2014; Racimo *et al*. 2015). Therefore, haplotype-based methods may be sensitive to local inaccuracies in the estimation of recombination rate along the genome. Furthermore, although the magnitude of divergence was captured by the between-lineages spectrum of shared IBS tracts, the two within-lineages spectrums were highly similar, possibly leading to a lack of signal to precisely estimate the duration of secondary contact with the IBS tract method.

Finally, the two approaches implemented here neglect the effects of temporal changes in effective population size and selection against introgressed tracts. Populations surviving in glacial refugia possibly experienced bottlenecks (Hewitt 2000), which could have impacted the IBS tract spectrum (Harris and Nielsen 2013). Furthermore, the removal of blocks of foreign ancestry by selection may explain the over-predicted abundance of introgressed tracts shorter than 500 kb in the Mediterranean (Fig. 2) and of IBS tracts longer than 100 kb (Fig. 3). Future works will have to integrate the effect of selection against introgression within demographic models, as previously done for demographic inference from unphased data (Sousa *et al*. 2013; Roux *et al*. 2016; Rougeux *et al*. 2017).

### The remolding of genomics islands of differentiation

The role of gene flow in generating genomic islands is a long standing debate (Wu 2001; Cruickshank and Hahn 2014; Burri 2017a; Ravinet *et al*. 2017). In particular, whether gene flow simply remodels evolving or pre-existing divergence patterns or constrains divergence to evolve principally in low-recombining regions remains an open question (Yeaman *et al*. 2016). Our results demonstrate that linked selection (Cutter and Payseur 2013) has generated heterogeneous genome divergence between sea bass lineages during their geographic isolation. This was supported both by the genome-wide correlations between divergence and recombination and the reconstructed ancestral landscape of divergence. We previously showed that large-scale variation in recombination rate across the sea bass genome is responsible, in interaction with selection, for increased rates of lineage sorting in low-recombining regions (Tine *et al*. 2014). The reduction of recombination in the center of chromosomes relative to their peripheries is a common feature of fish genomes (Bradley *et al*. 2011; Roesti *et al*. 2012), which is possibly due to crossover interference (Allendorf *et al*. 2015) and increased recombination in telomeric regions during male meiosis (Lien *et al*. 2011). Therefore, linked selection generating heterogeneous differentiation across the genome can be seen as a null expectation for allopatric divergence in sea bass, as in other fish.

Gene flow after secondary contact has the potential to rapidly remodel heterogeneous divergence landscapes by eroding neutral differentiation (Barton and Bengtsson 1986; Yeaman *et al*. 2016). However, whether genomic islands generated by linked selection tend to contain reproductive isolation loci and to act preferentially as barriers to gene flow remains unresolved. Our direct detection of introgressed tracts revealed broad variation in the rate of introgression among regions displaying similar levels of divergence. In some regions where ancestral peaks of F_ST_ have been found, the amount of introgression was high and the peaks almost vanished after secondary contact. This suggests that these incidental islands do not contain reproductive isolation loci (Cruickshank and Hahn 2014). On the other hand, reduced introgression in many regions confirms the view that introgressed alleles generally have negative fitness effects in the foreign genetic background (Martin and Jiggins 2017), although we did not specifically address the extent of adaptive introgression. The spatial comparison of introgression patterns between recipient populations (W-MED and E-MED) at variable distances from the source population (ATL) provides information about the distribution of fitness effects of introgressed tracts. Some genomic islands were resistant to introgression in both Mediterranean populations, indicating strong selection against introgressed tracts. In most genomic islands, however, introgression was more reduced in the E-MED compared to the W-MED. This either suggests that stronger migration overwhelms the effect of selection in the W-MED, or that selection takes more time to remove weakly selected migrant tracts as they diffuse from the western to the eastern part of the Mediterranean Sea. An important result stemming from the analysis of introgression is that the degree of resistance to gene flow for a given region was positively related to the past level of differentiation and to the absolute divergence between haplotypes, independently of the local mutation rate. Therefore, the strength of selection against introgressed tracts at a given genomic location seems to be at least partly explained by the coalescence time between haplotypes.

The expected amount of absolute divergence in low-recombining regions of the sea bass genome can be determined by summing values of ancestral diversity (*θ*_anc_ ≈ 0.001, estimated from the mean diversity of contemporary populations) and sequence divergence due to the accumulation of mutations in both lineages after split (2*μT* ≈ 0.001, determined using the divergence time estimated from demographic models). This amounts to E(*d*_XY_) = *θ*_anc_ + 2*μT* = 0.002, a value twice higher than the observed genome-wide average, possibly due to introgression. By contrast, the strongest barriers to gene flow between Atlantic and Mediterranean sea bass involve regions with higher *d*_XY_ values ranging between 0.002 and 0.005. This excess of coalescence time is not expected under a scenario whereby differential introgression reshapes a heterogeneous divergence landscape previously established by linked selection alone (Burri 2017b). To explain these results, we thus need to consider the sorting of ancient polymorphisms during the divergence period, later acting as barrier loci upon secondary contact.

The origin of such alleles that started to diverge before the average coalescent time expected from historical reconstructions remains uncertain. Recent studies have emphasized the role of introgression from a distantly related lineage as a source of new adaptations (Dasmahapatra *et al*. 2012; Huerta-Sánchez *et al*. 2014; Lamichhaney *et al*. 2015), which can possibly play a role in reproductive isolation. Although we found no evidence for contemporary gene flow between *D. labrax* and its closest relative *D. punctatus* (Tine *et al*. 2014), we cannot rule out a possible past admixture with *D. punctatus* or an extinct lineage, or the existence of an ancestral population structure (Racimo *et al*. 2015). Another possible explanation involves the existence of balanced polymorphisms maintained for a long time in the ancestral population, followed by the fixation of alternative alleles in the derived populations (Guerrero and Hahn 2017).

Whatever the origin of the alleles that contribute to reproductive isolation, our results clearly show that barrier loci tend to map preferentially to low-recombining regions. Such pattern may arise through different but non-mutually exclusive mechanisms. First, during isolation, weakly deleterious mutations are more likely to become fixed by drift or due to hitchhiking with a positively selected allele if recombination is reduced (Birky and Walsh 1988). This may trigger the fixation of compensatory mutations independently in each population, which could become genetic incompatibilities upon contact. Another effect of linked selection is to accelerate lineage sorting (Cruickshank and Hahn 2014), increasing the chance to fix alternative alleles in low-recombining regions during short isolation periods (i.e. < 10 *N*_e_ generations). Finally, during secondary contacts, the retention of divergence is facilitated when multiple incompatibility loci combine their effects through linkage. Therefore, the density of selected sites determines, in interaction with recombination, the strength of selection against introgressed tracts and the tendency for increased neutral introgression near chromosome extremities (Barton and Bengtsson 1986; Yeaman *et al*. 2016; Aeschbacher *et al*. 2017; Martin and Jiggins 2017). These different effects are likely to be amplified if ancestral variation is fueled by foreign alleles coming from a distant lineage during the divergence history.

## Conclusion

To conclude, our results shed new light on the origin and remolding of genomic islands during the divergence history of European sea bass lineages. Thanks to the use of haplotypic information, we provide a more mechanistic understanding of the complex interplay between linked selection, allele age and resistance to introgression. The recombination landscape appears to be an essential driver of the observed genomic patterns, influencing both lineage sorting and introgression. Our findings also support that the genomic islands generated by linked selection tend to be disproportionately involved in reproductive isolation during allopatric speciation, although some of them are purely incidental and are currently being eroded by gene flow. However, revealing the broad range of selective effects associated with the divergent regions found in speciation genomics studies may necessitate moderate to high levels of contemporary gene flow. Finally, the probability of introgression in a particular genomic region is negatively related to the level of divergence between alleles, a result possibly indicating that either past admixture or long-term balancing selection has participated to the evolution of reproductive barriers in sea bass.

## References

Abbott R., Albach D., Ansell S., Arntzen J. W., Baird S. J. E., et al., 2013 Hybridization and speciation. J. Evol. Biol. 26: 229–246.

Aeschbacher S., Selby J. P., Willis J. H., Coop G., 2017 Population-genomic inference of the strength and timing of selection against gene flow. PNAS 114: 7061–7066.

Allendorf F. W., Bassham S., Cresko W. A., Limborg M. T., Seeb L. W., et al., 2015 Effects of Crossovers Between Homeologs on Inheritance and Population Genomics in Polyploid-Derived Salmonid Fishes. J Hered 106: 217–227.

Assefa S., Keane T. M., Otto T. D., Newbold C., Berriman M., 2009 ABACAS: algorithm-based automatic contiguation of assembled sequences. Bioinformatics 25: 1968–1969.

Barton N., 1979 Gene flow past a cline. Heredity 43: 333–339.

Barton N., Bengtsson B. O., 1986 The barrier to genetic exchange between hybridising populations. Heredity 57: 357–376.

Birky C. W., Walsh J. B., 1988 Effects of linkage on rates of molecular evolution. PNAS 85: 6414–6418.

Bradley K. M., Breyer J. P., Melville D. B., Broman K. W., Knapik E. W., et al., 2011 An SNP-Based Linkage Map for Zebrafish Reveals Sex Determination Loci. G3: Genes, Genomes, Genetics 1: 3–9.

Browning S. R., Browning B. L., 2011 Haplotype phasing: existing methods and new developments. Nat Rev Genet 12: 703–714.

Burri R., 2017a Interpreting differentiation landscapes in the light of long-term linked selection. Evolution Letters.

Burri R., 2017b Dissecting differentiation landscapes: a linked selection’s perspective. J. Evol. Biol. 30: 1501–1505.

Charlesworth B., Morgan M. T., Charlesworth D., 1993 The effect of deleterious mutations on neutral molecular variation. Genetics 134: 1289–1303.

Coyne J. A., Orr A. H., 2004 Speciation. Sunderland MA, massachusetts U.S.A.

Cruickshank T. E., Hahn M. W., 2014 Reanalysis suggests that genomic islands of speciation are due to reduced diversity, not reduced gene flow. Mol Ecol 23: 3133–3157.

Cutter A. D., Payseur B. A., 2013 Genomic signatures of selection at linked sites: unifying the disparity among species. Nat Rev Genet 14: 262–274.

Danecek P., Auton A., Abecasis G., Albers C. A., Banks E., et al., 2011 The variant call format and VCFtools. Bioinformatics 27: 2156–2158.

Darling A. C. E., Mau B., Blattner F. R., Perna N. T., 2004 Mauve: Multiple Alignment of Conserved Genomic Sequence With Rearrangements. Genome Res. 14: 1394–1403.

Dasmahapatra K. K., Walters J. R., Briscoe A. D., Davey J. W., Whibley A., et al., 2012 Butterfly genome reveals promiscuous exchange of mimicry adaptations among species. Nature 487: 94–98.

Ellegren H., Smeds L., Burri R., Olason P. I., Backström N., et al., 2012 The genomic landscape of species divergence in Ficedula flycatchers. Nature 491: 756–760.

Feder J. L., Egan S. P., Nosil P., 2012 The genomics of speciation-with-gene-flow. Trends in Genetics 28: 342–350.

Gagnaire P.-A., Pavey S. A., Normandeau E., Bernatchez L., 2013 The Genetic Architecture of Reproductive Isolation During Speciation-with-Gene-Flow in Lake Whitefish Species Pairs Assessed by Rad Sequencing. Evolution 67: 2483–2497.

Guerrero R. F., Hahn M. W., 2017 Speciation as a Sieve for Ancestral Polymorphism. Mol Ecol.

Harr B., 2006 Genomic islands of differentiation between house mouse subspecies. Genome Res. 16: 730–737.

Harris K., Nielsen R., 2013 Inferring Demographic History from a Spectrum of Shared Haplotype Lengths. PLOS Genet 9.

Harrison R. G., Larson E. L., 2016 Heterogeneous genome divergence, differential introgression, and the origin and structure of hybrid zones. Mol Ecol 25: 2454–2466.

Hewitt G. M., 1988 Hybrid zones-natural laboratories for evolutionary studies. Trends in Ecology & Evolution 3: 158–167.

Hewitt G. M., 1996 Some genetic consequences of ice ages, and their role in divergence and speciation. Biological Journal of the Linnean Society 58: 247–276.

Hewitt G., 2000 The genetic legacy of the Quaternary ice ages. Nature 405: 907–913.

Huerta-Sánchez E., Jin X., Asan, Bianba Z., Peter B. M., et al., 2014 Altitude adaptation in Tibetans caused by introgression of Denisovan-like DNA. Nature 512: 194–197.

Kelleher J., Etheridge A. M., McVean G., 2016 Efficient Coalescent Simulation and Genealogical Analysis for Large Sample Sizes. PLOS Comput Biol 12: e1004842.

Lamichhaney S., Berglund J., Almén M. S., Maqbool K., Grabherr M., et al., 2015 Evolution of Darwin’s finches and their beaks revealed by genome sequencing. Nature 518: 371–375.

Lawson D. J., Hellenthal G., Myers S., Falush D., 2012 Inference of Population Structure using Dense Haplotype Data. PLOS Genet 8.

Lemaire C., Versini J.-J., Bonhomme F., 2005 Maintenance of genetic differentiation across a transition zone in the sea: discordance between nuclear and cytoplasmic markers. Journal of Evolutionary Biology 18: 70–80.

Li H., 2013 Aligning sequence reads, clone sequences and assembly contigs with BWA-MEM. arXiv: preprint.

Liang M., Nielsen R., 2014 The Lengths of Admixture Tracts. Genetics 197: 953–967.

Lien S., Gidskehaug L., Moen T., Hayes B. J., Berg P. R., et al., 2011 A dense SNP-based linkage map for Atlantic salmon (Salmo salar) reveals extended chromosome homeologies and striking differences in sex-specific recombination patterns. BMC Genomics 12: 615.

Marques D. A., Lucek K., Meier J. I., Mwaiko S., Wagner C. E., et al., 2016 Genomics of Rapid Incipient Speciation in Sympatric Threespine Stickleback. PLOS Genetics 12.

Martin S. H., Jiggins C. D., 2017 Interpreting the genomic landscape of introgression. Current Opinion in Genetics & Development 47: 69–74.

Maynard Smith J. M., Haigh J., 1974 The hitch-hiking effect of a favourable gene. Genetics Research 23: 23–35.

Mayr E., 1942 Systematics and the origin of species, from the viewpoint of a zoologist. Harvard University Press.

McKenna A., Hanna M., Banks E., Sivachenko A., Cibulskis K., et al., 2010 The Genome Analysis Toolkit: A MapReduce framework for analyzing next-generation DNA sequencing data. Genome Res. 20: 1297–1303.

Nachman M. W., Payseur B. A., 2012 Recombination rate variation and speciation: theoretical predictions and empirical results from rabbits and mice. Phil. Trans. R. Soc. B 367: 409–421.

Nadeau N. J., Whibley A., Jones R. T., Davey J. W., Dasmahapatra K. K., et al., 2012 Genomic islands of divergence in hybridizing Heliconius butterflies identified by large-scale targeted sequencing. Phil. Trans. R. Soc. B 367: 343–353.

Nosil P., Funk D. J., Ortiz-Barrientos D., 2009 Divergent selection and heterogeneous genomic divergence. Molecular Ecology 18: 375–402.

Pease J., Rosenzweig B., 2015 Encoding Data Using Biological Principles: the Multisample Variant Format for Phylogenomics and Population Genomics. IEEE/ACM Transactions on Computational Biology and Bioinformatics PP: 1–1.

Pool J. E., Nielsen R., 2009 Inference of Historical Changes in Migration Rate From the Lengths of Migrant Tracts. Genetics 181: 711–719.

Racimo F., Sankararaman S., Nielsen R., Huerta-Sánchez E., 2015 Evidence for archaic adaptive introgression in humans. Nat Rev Genet 16: 359–371.

Ravinet M., Faria R., Butlin R. K., Galindo J., Bierne N., et al., 2017 Interpreting the genomic landscape of speciation: a road map for finding barriers to gene flow. J. Evol. Biol. 30: 1450–1477.

Rissman A. I., Mau B., Biehl B. S., Darling A. E., Glasner J. D., et al., 2009 Reordering contigs of draft genomes using the Mauve Aligner. Bioinformatics 25: 2071–2073.

Roesti M., Hendry A. P., Salzburger W., Berner D., 2012 Genome divergence during evolutionary diversification as revealed in replicate lake–stream stickleback population pairs. Molecular Ecology 21: 2852–2862.

Rosenzweig B. K., Pease J. B., Besansky N. J., Hahn M. W., 2016 Powerful methods for detecting introgressed regions from population genomic data. Mol Ecol: 2387–2397.

Rougeux C., Bernatchez L., Gagnaire P.-A., 2017 Modeling the Multiple Facets of Speciation-with-Gene-Flow toward Inferring the Divergence History of Lake Whitefish Species Pairs (Coregonus clupeaformis). Genome Biol Evol 9: 2057–2074.

Roux C., Fraïsse C., Romiguier J., Anciaux Y., Galtier N., et al., 2016 Shedding Light on the Grey Zone of Speciation along a Continuum of Genomic Divergence. PLOS Biology 14.

Sedghifar A., Brandvain Y., Ralph P., 2016 Beyond clines: lineages and haplotype blocks in hybrid zones. Mol Ecol 25: 2559–2576.

Seehausen O., Butlin R. K., Keller I., Wagner C. E., Boughman J. W., et al., 2014 Genomics and the origin of species. Nat Rev Genet 15: 176–192.

Snyder C. W., 2016 Evolution of global temperature over the past two million years. Nature 538: 226–228.

Sousa V. C., Carneiro M., Ferrand N., Hey J., 2013 Identifying Loci Under Selection Against Gene Flow in Isolation-with-Migration Models. Genetics 194: 211–233.

Tine M., Kuhl H., Gagnaire P.-A., Louro B., Desmarais E., et al., 2014 European sea bass genome and its variation provide insights into adaptation to euryhalinity and speciation. Nature Communications 5: 5770.

Turner T. L., Hahn M. W., Nuzhdin S. V., 2005 Genomic Islands of Speciation in Anopheles gambiae. PLOS Biol 3.

Van der Auwera G. A., Carneiro M. O., Hartl C., Poplin R., Angel G. del, et al., 2013 From FastQ data to high confidence variant calls: the Genome Analysis Toolkit best practices pipeline. Curr Protoc Bioinformatics 11: 11–10.

Wolf J. B. W., Ellegren H., 2016 Making sense of genomic islands of differentiation in light of speciation. Nat Rev Genet 18: 87–100.

Wu C.-I., 2001 The genic view of the process of speciation. Journal of Evolutionary Biology 14: 851–865.

Yeaman S., Aeschbacher S., Bürger R., 2016 The evolution of genomic islands by increased establishment probability of linked alleles. Mol Ecol 25: 2542–2558.

